# Genomic Structural Variation Rescues a Classic Biological Invader from a Population Bottleneck

**DOI:** 10.64898/2026.01.30.702330

**Authors:** Christopher A. Osborne, Brian M. Foote, Steven J. Fleck, Hannah M. Waterman, Sarah L. Chang, Melia G. Nafus, M. Renee Bellinger, Levi N. Gray, Trevor J. Krabbenhoft

**Affiliations:** Department of Biological Sciences, University at Buffalo, Buffalo, NY, USA; U.S. Geological Survey, Pacific Island Ecosystems Research Center, Hawaii National Park, Hawaii, HI 96718, USA

**Keywords:** Genetic diversity, evolvability, population viability, 50/500 Rule, Lewontin’s Paradox, structural variants, inbreeding, pangenome, conservation genetics, genetic rescue, Population-Scale Long Read Sequencing, Structural variant ROH

## Abstract

Invasion genetics presents a classic paradox: how do species successfully spread despite severe population bottlenecks? The brown treesnake (*Boiga irregularis*) in Guam represents a striking example of this phenomenon, having been introduced with only a handful of individuals. We show that the population endured an extreme bottleneck, with roughly half of the genome exhibiting runs of homozygosity, comparable to species of conservation concern. Despite this, we uncovered extensive diversity in the form of nearly 19,000 genomic structural variants, which affect almost eight times more of the genome than single-nucleotide variants and provide material for ‘rescuing’ the population from inbreeding-driven declines. Structural variant density was highest in gene promoters, where recombination and DNA repair often occur, providing a mechanism for rapid evolution of gene-linked diversity. This diversity is enriched in genes vital for adaptive immunity and olfaction, suggesting genomic diversity in key chromosomal regions can rescue populations from inbreeding. This work has critical implications for invasion biology and conservation genetics practitioners.

**TEASER:** Genomic structural variants rescue a textbook biological invader from a population bottleneck and inbreeding

## INTRODUCTION

A long-standing tenet of population genetics theory is that genetic diversity is positively correlated with population viability and evolvability across lineages (*1–3*). Central to this theory is the requirement of heritable genetic variation in a trait, upon which natural selection can act (*4*). Reductions in genetic diversity, therefore, may reduce the ability of a population to adapt to novel environmental conditions (*5*, *6*). This logic has been widely applied in invasion biology and conservation genomics, which posit that populations with reduced genetic diversity (and concomitantly, reduced evolvability) are at greater risk of extinction or extirpation (*7–11*). While there is a large amount of theoretical and empirical evidence to support this theory (*12*, *13*), a growing body of literature suggests the relationship between traditional metrics for genetic diversity and population viability/adaptability is more nuanced than previously thought (*14–17*). For example, there are several instances of populations with effectively no genetic diversity (i.e., nucleotide diversity or heterozygosity at measured sites) that do not seem to suffer reduced fitness (*18–24*), sometimes even having lower levels of genetic load than larger populations (*17*). Paradoxically, there are now many documented instances of highly inbred populations exhibiting rapid and complex adaptive evolution (*21*, *25–28*).

One potential explanation for the disconnect between traditional metrics of genetic diversity and adaptability or population viability may stem from an overreliance on measures of nucleotide-based population genetic diversity metrics as indicators of “genetic health” (*16*, *29*). Recent work has shown that neutral and functional genetic diversity are often poorly correlated (*15*, *30*), such that functional diversity is likely a far more informative measure of a population’s adaptive potential (*31*). For instance, Mather et al. (*32*) found that while neutral and functional genetic diversity correlate in an endangered rattlesnake, neutral markers failed to predict adaptive potential in nonequilibrium populations reliably. Similarly, Anstett et al. (*33*) demonstrated that only levels of genetic diversity at a handful of adaptive loci, but not at many neutral loci, were predictive of natural evolutionary rescue in the scarlet monkeyflower, *Mimulus cardinalis*. Implicit in these findings is that different regions of the genome respond heterogeneously to evolutionary pressures.

Those studies highlight another potential source of disconnect between genetic diversity and adaptability: an over-reliance on genome-wide measures of genetic diversity when assessing populations. Using whole-genome summary metrics of genetic diversity is likely to miss local regions of chromosomes that may play a disproportionate role in organismal fitness (*34*, *35*). Genome-wide measures of diversity, such as heterozygosity, effective population size (*N_e_*), and inbreeding coefficients like the fraction of the genome in runs of homozygosity (fROH), are therefore likely to provide an incomplete picture of the adaptive potential, especially when the sole focus is at the level of nucleotide variation. This argument is bolstered by a growing appreciation of the importance of “islands of divergence”, which highlights that different regions of the genome evolve differently and, therefore, may differ in their evolutionary roles (*36–39*). As such, investigating more locus-specific patterns of genetic diversity may provide a more accurate assessment of the “genetic health” or evolutionary potential of populations.

High-quality genome assemblies and long-read sequencing have revealed the pervasiveness and evolutionary significance of genomic structural variants (SV; mutations affecting ≥50 base pairs (*40–43*). Indeed, incorporating SVs into analyses attempting to link genotypes-to-phenotypes has been shown to drastically reduce the missing heritability phenomenon (*43–45*). Despite their well-documented importance, few studies investigating the connection between genetic diversity and population viability account for the presence and putative impact of SVs (*46*). This practice is a potentially consequential omission, as two common sources of SVs, transposable element activity (*47*, *48*), and gene copy number variants (*49*, *50*) have been associated with generating *de novo* mutations putatively linked to adaptation in highly inbred populations. Incorporating analyses capable of detecting SVs and functional genomic variation, therefore, represents an opportunity to improve population and conservation genetics research and better integrate different types of genetic diversity with their contributions to population viability (*46*, *51*, *52*).

Biological invasions present an opportunity to investigate the contribution of different types of genetic variation (single-nucleotide variations, SVs, and copy number variants) to population viability. Paradoxically, many successful invasive species establish, spread, and outcompete native species, sometimes through rapid adaptation, despite low initial levels of genetic diversity as measured by traditional metrics (*53–56*). We refer to the apparent decoupling of genetic diversity and population viability as the Genetic Diversity-Population Viability Paradox, “GD-PV Paradox”. While phenotypic, ecological, and behavioral factors undoubtedly play an essential role in invasion success (*57*, *58*), rapid adaptive evolution often underlies biological invasions (*53*, *55*, *59*). Further investigation into the genomic consequences of founding events and the genetic basis for invasion success will likely strengthen our understanding of the type of genetic diversity required for adaptive evolution.

The brown treesnake (*Boiga irregularis*) is a famous example of a biological invader driving extinctions and dramatically altering biodiversity through direct negative impacts on naïve prey items, disruption of food webs, and significantly altering large scale ecosystem function. This species represents a striking example of a highly successful invasive species that exhibits remarkably low levels of genetic diversity. Native to eastern Indonesia, Papua New Guinea, the Solomon Islands, and northern and eastern Australia, *B. irregularis* established and spread across the island of Guam, causing widespread extinction and cascading ecological perturbations including loss or reduction of several endemic bird species after it was unintentionally introduced in the mid-20^th^ century (*60–62*). However, prior genetic assessments using reduced representation sequencing methods suggested that the invasion success of *B. irregularis* occurred despite evidence of considerable inbreeding, consequent to a founding event from a single source population of 10 or fewer individuals originating from Manus Island (*63*). Furthermore, the degree of inbreeding did not appear to significantly affect the mating or reproductive success of *B. irregularis* (*64*). These findings have been interpreted as *B. irregularis*’ invasion success being driven by ecological and behavioral plasticity rather than by genetic adaptation. The *B. irregularis* invasion presents an ideal opportunity to test the GD-PV Paradox by examining the genomic consequences of the founding event. Here we offer a potential resolution of this paradox by offering a mechanistic explanation for this species’ invasion success in Guam.

We generated a haplotype-resolved chromosome-level genome assembly for *B. irregularis* from its recently expanded range using newly developed deep learning-based read correction and genome assembly methods. This approach allowed us to assemble the Z and W chromosomes independently and subsequently explore sex chromosome evolution in this species for the first time, and how it relates to the Guam invasion. Long-read genome resequencing data for eight additional *B. irregularis* samples enabled us to investigate genomic diversity (SNPs and SVs) at whole-genome and locus-specific scales, revealing limitations of using traditional whole-genome summary metrics to assess genomic diversity in a vertebrate that recently experienced an extreme population bottleneck. We provide genetic evidence that helps resolve the GD-PV Paradox and highlights how a broader perspective on the types of genetic diversity that can rescue inbred populations. This perspective is relevant, as a massive uptick in “genetic rescue” studies is occurring, but these studies rarely incorporate the types of genetic diversity evaluation that we show here to influence population viability.

## RESULTS

### Genome Assembly and Haplotype Phasing

We produced a high-quality reference genome for *B. irregularis* by generating 74.5 Gb (∼42.6x coverage) of Q20+ Oxford Nanopore Technologies long read sequences, as well as 46 Gb (∼27x coverage) high-throughput chromatin conformation capture (Hi-C) data from an adult female collected from Cocos Island, a barrier islet located just 2 km off the southern tip of Guam, located in the Mariana Islands. HERRO-corrected nanopore reads, combined with Hi-C data and assembled with Hifiasm, produced two highly-contiguous haplotype-resolved assemblies with contig N50s of 45.5 Mb and 37.9 Mb for haplotypes 1 and 2, respectively. Contigs from both assemblies were corrected and scaffolded using Hi-C data, producing final assemblies with scaffold N50s of 220 and 149 Mb and 97.3% and 94.6% of the assemblies anchored onto 18 chromosomes for haplotypes 1 and 2, respectively. Although the karyotype of *B. irregularis* is unknown, four other *Boiga* species have 18-chromosome karyotypes (*65*, *66*), suggesting that our scaffolding likely represents the true karyotype of this species. Hereafter, we refer to haplotype 1 as hapW and haplotype 2 as hapZ, as they contain the W and Z sex chromosomes, respectively (refer to the section on sex chromosome identification below). Haplotype-phased blocks generated for the remaining chromosomes (i.e., autosomes) were randomly assigned to either “haplotype”. We refer to these assemblies as hapW and hapZ solely to indicate which sex chromosome accompanies each randomized autosomal set.

HapW is shorter in length (1.65 Gb) than hapZ (1.91 Gb). While low heterozygosity between haplotypes can lead to erroneous variation in genome assembly sizes for haplotype-resolved assemblies (*67*, *68*), we provide evidence that the length difference between our assemblies is mainly due to differences in sex chromosomes and not poor haplotype phasing. For example, except for the Z and W chromosomes, homologous chromosomes only varied in length by 2.67% on average between haplotypes, and each chromosome had similar telomeric sequence density and locations (**Fig. 1b**). Additionally, the total length of telomeric repeats across autosomal chromosomes was comparable between hapZ (94.43 b) and hapW (80.94 Kb). Furthermore, while both haplotype assemblies had a high degree of completeness according to Benchmark Universal Single Copy Orthologs (BUSCO) analysis, hapZ had a noticeably higher completeness score (97.3%, 60 missing genes) than hapW (92.3%, 233 missing genes). The discrepancy in BUSCO scores is entirely driven by differences in the Z and W chromosomes, as the BUSCO score of hapZ is 92.2% with 235 missing BUSCO genes when the Z chromosome is removed from the assembly (**Fig. 1c**). Finally, the number of scaffolding gaps in each assembly was very similar between haplotypes and was in the single digits for all but four chromosomes (**Fig. 1e**). Collectively, this provides strong evidence of high assembly completeness of both haplotypes.

**Fig. 1.**
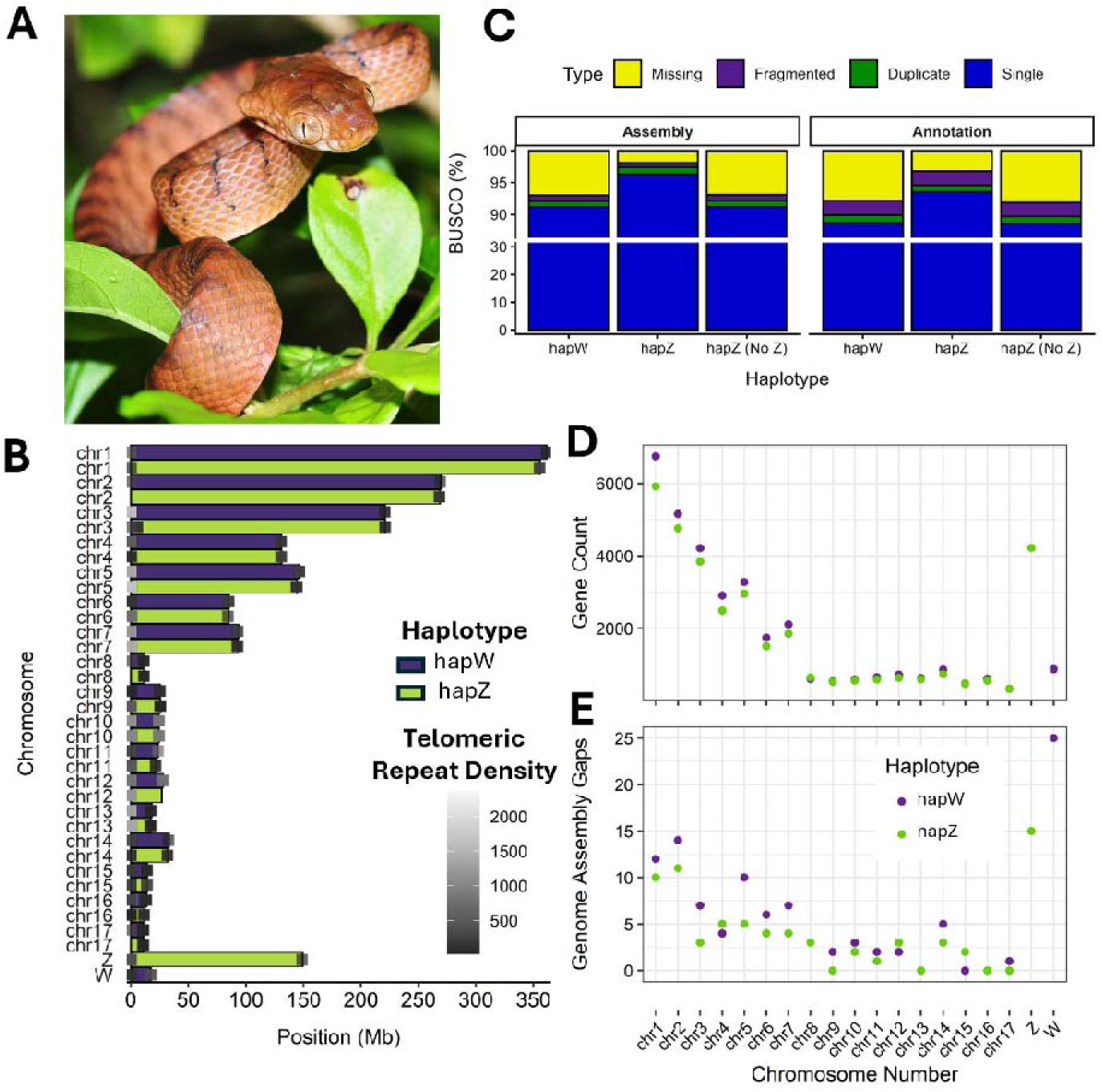
Genome Assembly and Haplotype Phasing for *Boiga irregularis*. **A**) Photograph of *B. irregularis* on Guam, photo credit Pavel Kirilliov CC BY-SA 2.0, https://creativecommons.org/licenses/by-sa/2.0, via Wikimedia Commons. **B**) Chromosome ideogram showing the full length of each chromosome in each haplotype. Boxes represent the region of each chromosome where the highest telomeric repeat (AACCCT) densities (per 20 Kb window) were detected. Boxes are shaded according to the total number of consecutive telomeric repeats found in each region. **C**) Benchmark Universal Single Copy Orthologs (BUSCO) scores for hapW and hapZ, and hapZ excluding the Z chromosome, for the *B. irregularis* genome assembly and annotations produced in this study. **D, E**) Gene count and number of genome assembly gaps found in each chromosome from the hapW and hapZ assemblies.

*Ab initio* and homology-based genome annotation tools using protein evidence from 16 other snake species (**Supplementary Table 1**) yielded 37,599 and 42,692 gene models, representing 89.9% and 94.6% of Vertebrata BUSCO genes for hapW and hapZ, respectively. The discrepancy in BUSCO scores between haplotypes was again driven entirely by sex chromosome dimorphism. The complete BUSCO score for the hapZ was 89.7% when gene models from the Z chromosome were excluded (**Fig. 1c**), and the number of annotated genes was similar on homologous autosomes across haplotypes (**Fig. 1d)**, providing strong evidence that both haplotypes were well annotated.

### Long-Term and Recent Demography

We investigated the long-term demographic history of *B. irregularis*, by generating approximately 20x genome coverage of Illumina NovaSeq 6000 short-read sequencing (SRS) data from the same individual used to produce the genome assembly and used these data to reconstruct variation in effective population size (*N_e_*) through time using Pairwise Sequentially Markovian Coalescence (PSMC [(*69*)]; **Fig. 2a**). To ensure that our PSMC results did not include an artifactual *N_e_* peak that is common in recently bottlenecked populations, we split the first time window using methods described by (*70*) (**Supplementary Fig. 1**). Demographic history reconstruction revealed that *B. irregularis* reached its largest estimated *N_e_* of ∼200,000 approximately 250 thousand years ago (Kya), followed by a rapid decline in *N_e_*, reaching a low point of ∼20,000 around 40 Kya before rising again to ∼60,000 until 3 Kya, after which it experienced a precipitous drop to an estimated *N_e_* of 980.

**Fig. 2.**
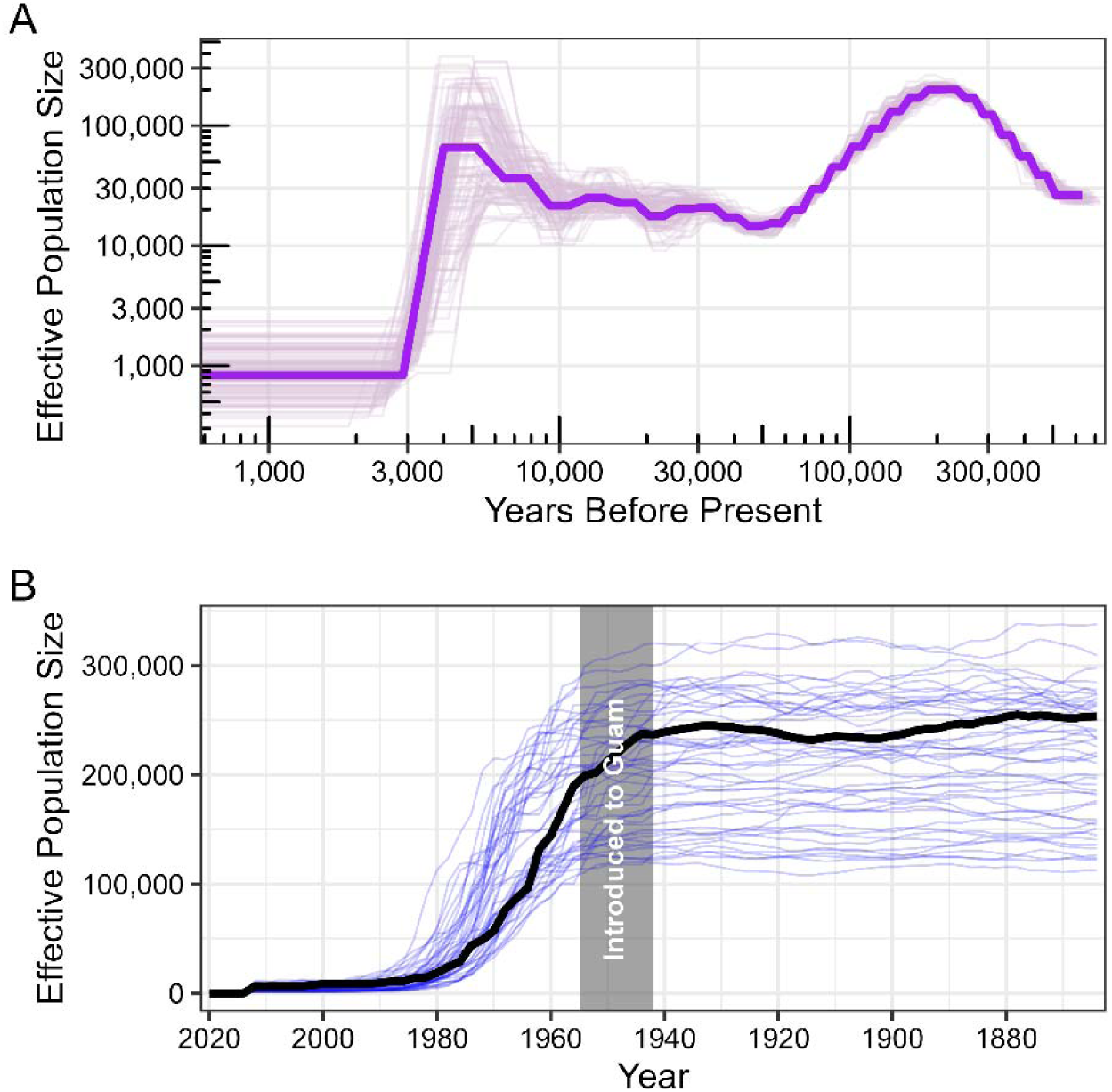
*Boiga irregularis* Experienced a Severe Population Bottleneck Following its Introduction to Guam. **A)** Long-term changes in the demographic history of *B. irregularis* estimated using the pairwise sequentially Markovian coalescent model (PSMC; [*69*]). A generation time of five years was used according to (*75*) and a mutation rate of µ = 7.29e-09 (*76*). Faint purple lines represent the results of 100 bootstrap replicates. **B)** Effective population size estimates for *B. irregularis* using the linkage disequilibrium-based tool GONE2 (*72*). Faint blue lines represent the results of 50 bootstrap replicates. The gray box denotes the approximate timing of *B. irregularis*’ introduction to Guam, between years 1942 and 1955 (*63*).

Markovian coalescence-based estimates of *N_e_* have been shown to perform poorly for estimates more recent than 10 Kya (*71*) and therefore, provide little insight into the demographic characteristics of *B. irregularis* following its introduction to Guam around 1945. We generated moderate coverage long-read resequencing data from eight additional *B. irregularis* samples (n = 3 females and n = 5 males) collected from Guam to estimate more recent (i.e., < 70 generations) changes in *N_e_*size using GONE2 (*72*). After quality filtering, excluding SNVs from the Z chromosome because sex chromosomes have unique inheritance patterns and sex-specific recombination rates compared to autosomes, and excluding single-nucleotide variants (SNVs) on any chromosomes shorter than 20 Mb (the minimum length threshold for this tool (*72*)), we were left with 1,073,398 SNVs across 11 autosomes for demographic analysis. *Boiga irregularis* began to experience a large reduction in *N_e_* from around 200,000 starting around 1954, which is within the estimated window of time for when it was introduced to Guam (1942–1955), and that this decline in *N_e_* continued until it reached around 59 in 2012, where it has remained until present (**Fig. 2b**). Alternatively, the bottleneck could have started earlier, such as during dispersal from Papua New Guinea to Manus Island prior to the introduction on Guam. Sequential island hopping via vegetation rafting or other means (*73*) could have led to progressive loss of diversity through small effective population sizes, although this is only weakly supported by the PSMC results. Regardless of the precise timing, *B. irregularis* was severely bottlenecked in Guam during the invasion.

### The *Boiga irregularis* Genome is Genetically Bottlenecked

The number and length of runs of homozygosity (ROH, segments of the genome that are inherited as nearly identical copies from both parents) are strong predictors of recent inbreeding events (*77*, *78*), and the genomic proportion contained within ROH (fROH) is highly predictive of inbreeding levels obtained from actual pedigree information (*79–81*). We identified the prevalence and length of ROH across the 17 autosomes of *B. irregularis*, to investigate patterns and extent of inbreeding in the Guam population. This analysis revealed a considerable amount of inbreeding as, on average, 49.2% (range = 42.8% to 63.1%) of the *B. irregularis* genome was contained within ROH (**Fig. 3a**). The average length of ROH ranged widely among samples, from 0.53 to 2.6 Mb, as did the length of the largest ROH, which ranged from 7.67 to 143 Mb (**Fig. 3a, Supplementary Table 2**). While shorter ROH (50 Kb – 0.99 Mb) were most common across all samples, all samples also contained at least four large ROH (>5 Mb), suggesting that both historical and recent inbreeding events have affected this population (**Fig. 3a**). Despite the high degree of inbreeding in each sample, ROH deserts (ROH detected in three or fewer samples) made up a larger portion of the genome than ROH islands (ROH detected in at least seven samples). Specifically, ROH deserts covered 453.7 Mb (30.4% of the genome) and contained 9,892 genes—nearly twice as many as the 5,964 genes found within the 297.7 Mb (20.0%) covered by ROH islands. Gene ontology (GO) enrichment analysis revealed that genes in ROH deserts were significantly enriched for several immune system terms, including major histocompatibility complex (MHC) genes, stress response, and olfactory-related GO functions (**Supplementary Data S1**). Genes in ROH islands were only significantly enriched for a single GO term, “cytoplasm” (**Supplementary Data S1**).

**Fig. 3.**
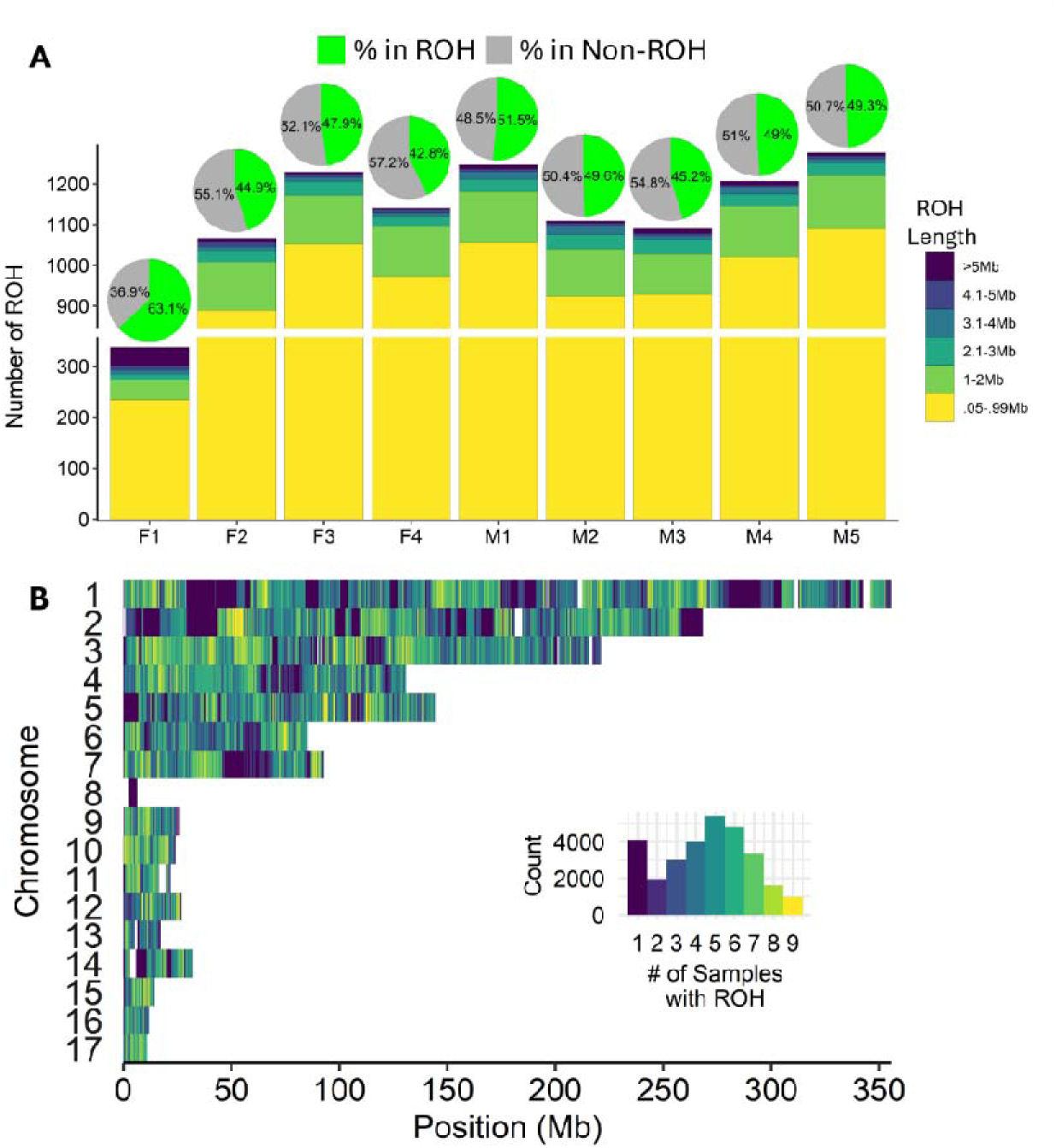
Extreme Runs of Homozygosity (ROH) are Present Across Much of the Genome in Boiga irregularis. **A)** ROH characteristics across four female (F1 - F4) and five male (M1 - M5) *B. irregularis* individuals. Pie charts depict the percentage of the genome contained within ROH (versus not) for each sample. The bar chart depicts the number and length distribution of ROH. **B)** ROH are widespread across *B. irregularis* genomes. Chromosomes are colored according to the number of samples for which a ROH was present in each window. The histogram insert depicts the color gradient and frequency of occurrence of overlapping ROH. ROH are present in all seventeen autosomes and are often shared by multiple individuals.

### Structural Variants Rescue Loss of Genetic Diversity from Inbreeding

To explore how genomic structural variants contribute to genetic diversity in *B. irregularis*, we implemented a read-mapping-based SV-calling pipeline. We used the nanopore reads from the eight resequenced samples, as well as the individual used to make the reference assembly, and detected 19,498 SVs. We then used an SV-genotyper (SVJedi-Graph v1.2.1 (*82*) to validate the accuracy of SV calls for each sample. The genotyper first constructed a *B. irregularis* variation graph using the newly detected SVs, then implemented read-to-graph alignments of the nanopore reads from each sample to produce a new set of SV calls. SVs with matching genotypes in both methods were considered validated. The variation graph revealed a high degree of SV calling accuracy, as 18,871 (96.8%) SVs were validated using this approach. Only SVs validated using the variation graph were used for further analysis. The final high-confidence SV set contained 10,169 deletions, 8,691 insertions, four duplications, and seven inversions (**Fig. 4a**).

**Fig. 4.**
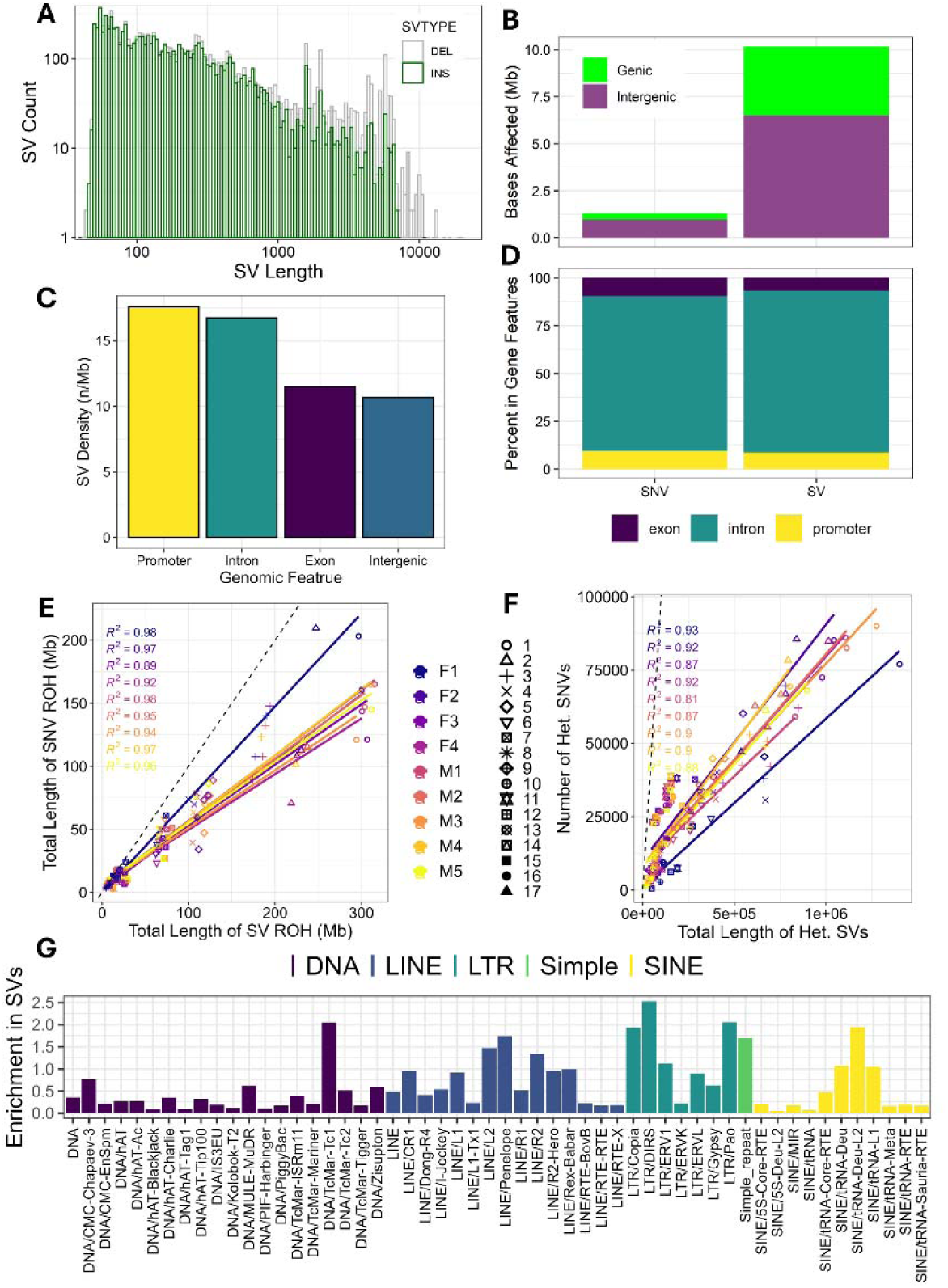
Structural Variants Represent a Significant Source of Genetic Diversity in the Inbred *Boiga irregularis*. **A)** Histogram displaying the length distribution of deletion and insertion structural variants (SVs) detected in nine *B. irregularis* from Guam. **B)** Bar chart showing the relative proportions of the genome affected by single-nucleotide variants (SNVs) and structural variants (SVs). **C)** Density of SVs across genomic features. The bar chart illustrates the frequency of SVs normalized per megabase (Mb) for four genomic subregions. **D)** Bar chart depicting the percentage of genic SNVs and SVs found in exons, introns, and promoter regions. **E)** Scatter plo comparing the total length of Runs of Homozygosity (ROH) calculated using SNVs and SV in megabases (Mb). **F)** Scatter plot comparing the total number of heterozygous SNVs and the total length of heterozygous SVs. For E and F, distinct markers represent individual chromosomes (1–17), and point colors correspond to sample IDs. The colored solid lines indicate linear regressions for each sample, with the corresponding R^2^ values shown. The black dashed line represents a 1:1 ratio for reference. **G)** Bar chart displaying fold-enrichment of TEs in SVs versus across the entire genome. Bars are colored by repeat class.

In total, SVs affected 10,191,284 bp of the genome, which is 7.9x more of the genome than was affected by SNVs, which only affected 1,295,475 bp (**Fig. 4b**). A similar discrepancy was observed for polymorphisms within genes. While far more genes were found to contain SNVs (n = 14,643 genes) versus SVs (n = 3,983 genes), SNVs only affected 344,596 bp (26.6% of SNVs) of gene sequence while SVs affected 3,661,413 bp (36% of SV-affected bases; **Fig. 4b**). Most SNVs and SVs affecting genes were found in introns (81.1% and 84%, respectively), and a larger portion of SNVs was found within exons (9.5%) and putative promoters (9.4%) than was the case for SVs (6.7% and 8.7%, respectively; **Fig. 4d**). Notably, SV density (SVs per megabase) was highest in putative gene promoters (17.58/Mb), followed by introns (16.74/Mb) and exons (11.5/Mb) (**Fig. 4c**). SV density was lowest in intergenic regions (10.66/Mb) despite these regions comprising ∼64% of the total sequence length. Genes affected by SVs were significantly enriched for olfactory-related GO functions, specifically “detection of chemical stimulus involved in sensory perception of smell”, “olfactory receptor activity”, and “sensory perception of smell” relative to all annotated genes (**Supplementary Fig. 2**).

The same biological features that make certain genomic regions SV hotspots—such as dense tandem arrays, local intergenic repeats, and elevated heterozygosity—also make them notoriously difficult to assemble properly. A key example is the major histocompatibility complex (MHC), an evolutionary hotspot that frequently resists contiguous assembly, even with long-read sequences (*83–85*). Indeed, we found that SVs were prevalent in MHC genes in our dataset (**Supplementary Data S2**). To investigate whether assembly artifacts impacted the SVs identified in this study, we assessed the assembly quality of the MHC locus via comparative mapping to the highly contiguous genome of the green cat snake (*B. cyanea*; (*86*), revealing that our original assembly was artificially fragmented into 11 contigs during the Hi-C scaffolding process (**Supplementary Fig. 3a**). After applying localized, reference-guided scaffolding to reconstruct a highly contiguous MHC in *B. irregularis* (refer to Methods section for details), we re-called SVs on the improved chromosome and confirmed that the region is indeed a genuine SV hotspot in *B. irregularis* (**Supplementary Fig. 3b-e**).

To evaluate the impact of SVs on genome-wide heterozygosity, we established a novel metric, “SV ROH”. Consistent with the minimum window size used for SNV ROH detection, we defined an SV ROH as a genomic tract of 50 Kb or longer containing zero heterozygous SVs. As expected, we observed a strong positive correlation between the total length of SV and SNV ROH within each sample. However, for most chromosomes across the sampled individuals, the total length of SV ROH was approximately twice as large as that of SNV-based ROH (**Fig. 4e**). This indicates that SNVs are more uniformly dispersed across the genome, interrupting runs of homozygosity more frequently than SVs. Crucially, despite the longer tracts of structural homozygosity, the cumulative number of base pairs altered by heterozygous SVs was, on average, 10.2 times greater than the total number of bases affected by heterozygous SNVs within each sample (**Fig. 4f**), suggesting the SVs represent a substantial reservoir of genetic diversity in this population. A total of 10,282 SVs (54.5% of all SVs) overlapped or contained transposable elements (TEs). Of these, LINE elements were the most common TE class, comprising over 67% of SV-TE associations with L2 and CR1 repeats representing the most abundant individual repeat families (**Supplementary Fig. 4**). Two LTR repeat classes, DIRS and Pao, as well as TcMar-Tc1 repeats, were SV-enriched, with over two times higher occurrence within SVs than in the rest of the genome (**Fig. 4g**).

### Structural Variants Drove Degeneration of the W Chromosome in *Boiga irregularis*

A comparison of the two assembly haplotypes revealed that one chromosome pair varied in size by over 100 Mb, far more than the length variation in the other chromosome pairs. To investigate whether this length difference represents sex chromosome heteromorphism, we searched for conserved synteny between the *B. irregularis* haplotype-resolved assemblies and 16 other snake species, including those for which the sex chromosomes have been previously validated. The chromosome pair with extreme size variation between haplotypes had conserved synteny with previously identified sex chromosomes in other snakes, suggesting the larger chromosome represents the Z and the smaller chromosome the W (**Fig. 5a**).

**Fig. 5.**
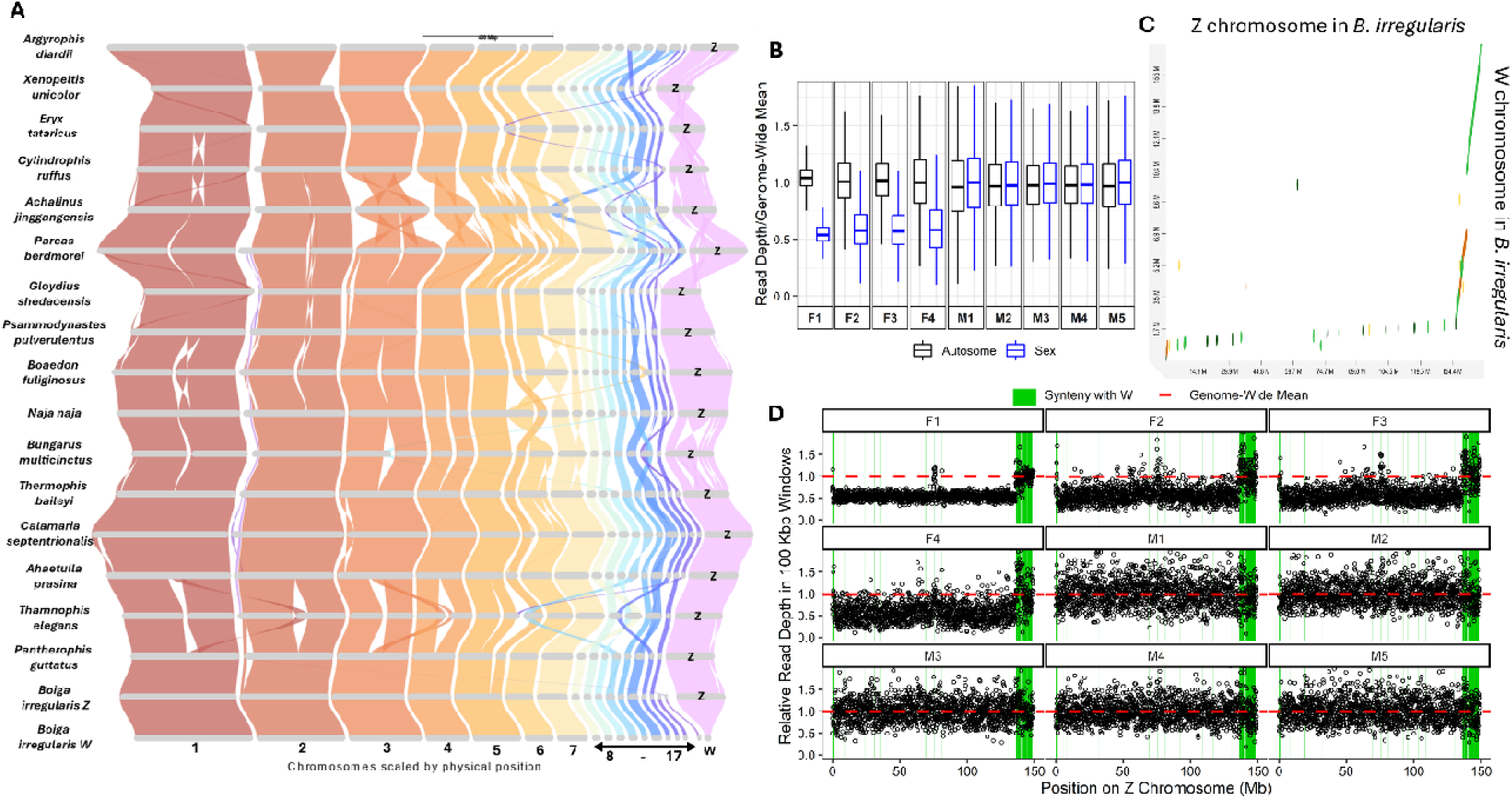
Sex Chromosome Heteromorphism in *Boiga irregularis*. **A)** Riparian plot illustrating conserved synteny between *B. irregularis* hapW and hapZ haplotype assemblies from this study and 16 other snake genomes from Peng et al. (*88*) (Supplemental Table 1). Bands are colored according to synteny with *B. irregularis* chromosomes. **B)** Box plot comparing the read depth of nanopore reads mapped to autosomes and sex chromosomes in *B. irregularis* male and female samples. **C)** Dot plot displaying the syntenic relationship between the Z and W chromosomes of *B. irregularis*. **D)** Relative read depth across the Z chromosome of *B. irregularis* (mean read depth calculated in non-overlapping 100 Kb windows divided by the genome-wide mean read depth for each sample). The red horizontal bar represents the genome-wide mean for each sample. Green vertical boxes correspond to regions of retained synteny between the Z and W chromosomes assembled for *B. irregularis* in this study.

We implemented a read-depth-based approach (*87*) to validate size differences in sex chromosomes to test if the significant disparity in size between the assembled Z and W chromosomes in *B. irregularis* noted above is the result of true heteromorphism or the product of assembly error. Briefly, the number of reads generated per locus should represent the number of copies of that locus in the genome, making read depth proportional to the copy number of that locus. Accordingly, the read depth at Z- or X-specific loci should be approximately half the genome-wide average in individuals of the heterogametic sex. This method, including PCR-based validation, was recently confirmed as a reliable approach for detecting sex chromosome-specific loci in a taxonomically diverse group of snakes (*87*). We mapped nanopore reads from the five male and four female samples to the newly assembled *B. irregularis* hapZ assembly (i.e., the assembly with the putative Z chromosome). Read depth for the putative Z chromosome was approximately half the average autosomal read depth in the female samples only (**Fig. 5a**). An approximately 17 Mb region of the Z chromosome had read depths matching the genome-wide average in female samples, possibly representing remnant pseudoautosomal or undegraded regions between the Z and W chromosomes (**Fig. 5b**). We mapped the assembled W chromosome to the Z chromosome, revealing that the regions with read depths matching the genome-wide average correspond to regions of strongly conserved synteny between the Z and W chromosomes (**Fig. 5c, d**), providing further evidence of a putative pseudoautosomal region.

These results validate our assembly and phasing results for the Z and W chromosomes, providing strong support for sex chromosome heteromorphism driven by structural variation-linked degeneration in *B. irregularis*.

## DISCUSSION

Genetic diversity is often low during successful biological invasions, providing a challenge to population genetics theory. We addressed this Genetic Diversity-Population Viability Paradox by producing a haplotype-resolved genome assembly and including genomic structural variants into our investigation of genetic diversity in brown treesnake in Guam. Our results provide a rich context for the genomic landscape of a highly successful invader and demonstrate that a focus solely on broad genetic diversity metrics may underestimate or misrepresent a population’s genetic diversity. We demonstrate that estimates of effective population size and inbreeding coefficients, which are among the most widely used metrics to assess the “genetic health” of populations, suggest that *B. irregularis* in Guam is genetically bottlenecked and exhibits a similar level of inbreeding as species of conservation concern. For example, the levels of inbreeding we detect in *B. irregularis* are similar to those found in the threatened eastern massasauga rattlesnake (*Sistrurus catenatus;* (*76*). However, unlike *B. irregularis,* which is currently comprised of a thriving population of around two million individuals (*75*, *89*), *S. catenatus* is estimated to have a few extant populations of less than 50 individuals (*90*) and is hypothesized to be experiencing population viability risks due to increased genetic load from inbreeding depression (*32*, *76*).

*Boiga irregularis*’ incredible success as an invasive species in Guam, despite exhibiting a similar degree of inbreeding as a threatened species suffering from inbreeding depression, adds to a growing list of empirical examples of genome-wide metrics of genetic diversity, especially those based on neutral diversity, poorly predicting a population or species’ viability or fitness (*14–17*). A likely reason for this discrepancy is that, despite roughly half of the *B. irregularis* genome being found in ROH, certain regions consistently retain heterozygosity among individuals. These regions—ROH deserts—tended to cluster around genes involved in immune processes, stress adaptation, and olfactory functions, pointing to a pattern where gene content drives localized genetic diversity. We also demonstrate that a considerable amount of genomic diversity, in the form of SVs, exists in this population. Genes affected by SVs were significantly enriched for the same olfaction-related GO terms as ROH deserts, suggesting that, in addition to SNVs, SVs may be playing a role in producing or maintaining diversity in these regions. Structural variants, which comprise nearly eight times more variation than SNVs alone (based on the number of nucleotides affected), are either not accounted for or, worse, skew the results of SNV-based genome-wide metrics, highlighting the importance of accounting for SVs when assessing the “genetic health” of a population.

The elevated genetic diversity observed in both major histocompatibility complex (MHC) and olfactory genes presents a putative mechanism explaining how genetic diversity was either maintained or re-established in this severely bottlenecked population. Elevated levels of genetic diversity in immune-related genes, especially in MHC genes, have been reported in otherwise highly inbred populations across a wide variety of taxa (*91–95*), including the aforementioned endangered eastern massasauga rattlesnake (*96*). One putative driver of MHC diversity in inbred populations is a type of sexual-selection-based balancing selection. Specifically, many vertebrates can use olfactory cues to select mates with genetically dissimilar or diverse MHC alleles (a form of disassortative mating), including several reptiles, which is theorized to provide a selective advantage by producing offspring with enhanced immune system breadth (*92*, *97–102*). The adaptive advantage of a more robust immune system is considered to be strong enough to maintain diversity, even in populations with small effective population sizes. Upon *B. irregularis’* introduction to Guam, exposure to a variety of pathogens to which its immune system would be relatively naive could have exerted strong selection pressure for the maintenance of a diverse adaptive immune system.

We also observed evidence of elevated genetic diversity in the olfaction genes putatively involved in detecting MHC-based pheromones. Vomeronasal receptors in the vomeronasal organ (VNO) are the primary mechanism by which vertebrates detect MHC diversity in their potential mates (*103*, *104*). In snakes, as in other squamates, the VNO is primarily composed of type-II vomeronasal receptors (V2R), which is likely the result of a shared gene family expansion in V2R genes (*105*). We functionally annotated 237 V2R genes in the *B. irregularis* genome, which were largely dispersed across multiple chromosomes, with the exception of a 60-copy cluster on the Z chromosome, a 63-copy cluster on chromosome 2, and a 23-copy cluster on chromosome 8 (**Supplementary Data S3**). Of these 237 V2R genes, 85 (35.8%) had at least one SV overlapping them, and 46 (19.4%) were contained within ROH deserts (**Supplementary Data S3**). The ratios of these estimates suggest diversity is being maintained or restored in these genes. It is possible that selection may be acting to maintain or regain diversity in key MHC genes, as well as the genes responsible for detecting MHC-based pheromone diversity in mates in the *B. irregularis* population in Guam. MHC-based mate preference has been documented in several reptiles (*106–109*), but future studies would be required to determine if this also occurs in *B. irregularis*. However, our discovery of two large V2R gene clusters, one on the Z chromosome, and another adjacent to the MHC locus on chromosome 2, provides support for a MHC-diversity-based mate preference system as a potential driver of genetic diversity in this population.

The elevated genetic diversity in select regions of the *B. irregularis* genome, despite high degrees of inbreeding in other regions, might explain why a previous study failed to detect an association between inbreeding levels and reproductive success in this population (*64*). If diversity were measured in and around MHC and V2R genes, an association between reproductive success and inbreeding might have been observed. Alternatively, chromosomal regions that harbor these hyperdiverse genes might be prone to both point mutations and structural variant occurrence due to the local chromatin structure, epigenetic state (e.g., DNA methylation), TE dynamics, and distance from the centromeres, all of which can affect recombination and mutation rates driving these features.

Our discovery that structural variants contribute nearly eight times more to genetic diversity than SNVs in the *B. irregularis* population in Guam directly supports the growing body of literature suggesting that standard, reference-based diversity metrics provide an incomplete picture of genome-wide polymorphisms (*51*, *52*, *110–112*). A recent meta-analysis in plants (*113*) argued that the weak scaling between population size and genetic diversity—a classic issue known as Lewontin’s Paradox (*114*)—may be partly explained by the failure of common SNV-based analyses to capture variation not well-represented in a reference genome. By using a reference-free, k-mer-based approach, Roberts and Josephs (*113*) found that diversity metrics scaled up to 20 times more strongly with population size proxies than traditional nucleotide diversity. Our findings in *B. irregularis* provide a clear example of their central hypothesis, as the thousands of previously uncharacterized SVs we identified represent the very type of non-SNV variation that their study suggests is systematically underestimated. This outcome indicates that considerable, unmeasured diversity present in complex genomic regions may be a critical, and often overlooked, factor in resolving the paradoxical relationship between a species’ census size and its molecular diversity.

These findings also serve as a cautionary tale for conservation genetics, where decisions about a species’ status and potential need for intervention often rely on nucleotide-based diversity metrics. Our results demonstrate that relying solely on SNV-based estimates can lead to an incomplete assessment of a population’s genetic health or viability. A population with low SNV diversity might be incorrectly deemed genetically impoverished and in need of “genetic rescue” when it could harbor substantial, unmeasured structural variation essential for its long-term survival and adaptability.

Structural variants in the *Boiga irregularis* genome had their highest density in gene promoter regions (versus introns, exons, and intergenic regions). In snakes, double-strand break repair and meiotic recombination occur at two distinct sets of sites: PRDM9 binding motifs and gene promoter regions (*115*). The high density of promoter-linked SVs in *B. irregularis* is consistent with a mechanism whereby double-strand break repair and meiotic recombination could produce hotspots of structural variation, leading to rapid evolution of gene-linked features in this inbred species. These findings have critical implications for understanding mechanisms of genetic rescue in bottlenecked populations. Future studies could explore the link between rapid evolution in SV hotspots and PRDM9 versus gene promoter-linked recombination in snakes.

Our comparison of SV- and SNV-based ROH highlights a critical, previously overlooked layer of genetic diversity in the *B. irregularis* population. If our analysis had relied exclusively on traditional SNV metrics, the full scope of genetic variation in this population would have been drastically underestimated. Because SNVs are more widely dispersed across the genome, they break up runs of homozygosity more frequently, resulting in shorter overall SNV ROH tracts compared to SV ROH. A superficial look at the expanded SV ROH lengths might suggest a heavily depleted genome; however, evaluating the actual footprint of these variants tells a different story. Heterozygous SVs affect over an order of magnitude (10.2x) more total base pairs than heterozygous SNVs. This finding demonstrates that while the *B. irregularis* population is highly inbred by traditional SNV standards, SVs maintain a massive, hidden reservoir of genomic variation. By altering large stretches of the genome, SVs introduce substantial structural and potentially functional diversity.

The precise mechanisms driving SV formation are complex and variable, but TEs often play a key role. We detected >2-fold enrichment of density of three classes of TEs in SVs versus the remainder of the genome, providing candidate drivers of SV formation for future studies. One of these classes, LTR-DIRS is often associated with centromere regions (*116*), where SVs are abundant, suggesting a potential role for particular TE classes in different areas on chromosomes. In summary, we show that SVs act as a primary driver of genetic diversity in this population, serving as a plausible mechanism of genetic “rescue” that provides the raw material necessary for the population’s continued adaptability and survival despite severe inbreeding.

Finally, using state-of-the-art haplotype-aware read-correction allowed us to fully assemble both *B. irregularis* sex chromosomes and demonstrate a role for structural variants in driving their differentiation. Although other haplotype-resolved assemblies for snakes exist (*117*, *118*), to the best of our knowledge, this represents the first time the sex chromosomes of a snake have been fully phased and assembled. Given that sex chromosomes in caenophidian snakes originated from autosomes (*119*, *120*), the independent assembly of the Z and W chromosomes allowed us to directly compare the degree of retained synteny between the sex chromosomes in this species, revealing that the W chromosome is only approximately 15% of the total length of the Z chromosome, a result we confirm with a read depth analysis. This finding suggests that the W chromosome has experienced considerable degeneration in *B. irregularis*, driven by structural variants. While other members of the Colubridae exhibit a range of sex chromosome heteromorphism, cytological evidence shows that at least two different species of *Boiga* have homomorphic sex chromosomes (*66*). *B. irregularis* is the only member in the genus with strongly heteromorphic sex chromosomes of those studied to date.

While the absence of data from the founding population on Manus Island prevents us from identifying the timing of this sex chromosome degeneration, one possibility is that the extreme inbreeding event experienced by *B. irregularis* may have facilitated the rapid degeneration of its W chromosome. It is possible that the W chromosome began degenerating with the Guam population founder event and subsequent population bottleneck, which could have severely decreased diversity in W chromosome alleles and been concurrently affected by relaxed selection. The natural lack of recombination on the W chromosome, coupled with the population bottleneck, likely compounded the effects of Muller’s Ratchet, where the chance loss of high-fitness genotypes leads to an irreversible ‘ratcheting’ of deleterious mutations throughout the population (*121*, *122*). The degenerated state of the W chromosome represents a series of large-scale structural variants that signify a permanent shift in the genomic architecture of *B. irregularis* in Guam - one that would have been missed using only genome-wide summary statistics. Future studies characterizing the level of sex chromosome heteromorphism in native populations of *B. irregularis* and other closely related congeners could provide insight into the timing of sex chromosome degeneration in this lineage.

Our use of read-mapping-based SV calling rather than conducting *de novo* assemblies for the eight resequenced individuals may have entailed technical trade-offs worth considering. For example, although SV detection methods that leverage alignments of assembled genomes have been shown to be superior for detecting larger SVs, especially large insertions, read-mapping-based approaches are better at detecting complex SVs such as translocations, inversions, and duplications (*123*). Read-mapping-based approaches also achieve higher genotyping accuracy when genome coverage is around 5 – 10x, as was the case with the resequencing data used in this study (*123*, *124*). As such, we may have failed to characterize some variants that would have been detected if we had used genome alignments for SV detection. Similarly, while it is possible that our moderate-coverage resequencing approach could have underestimated detection of SNVs and concomitantly overestimated ROH, our results show no correlation between fROH and average genome-wide sequencing coverage (**Supplementary Fig. 5**), suggesting that fROH estimates were not likely affected by variation in coverage. Another caveat is that ONT reads have higher basecalling error rates than PacBio Hifi reads, which can affect genome alignment and SNV detection accuracy. However, our nanopore reads had an average q-score of 23 (∼99.5% basecalling accuracy), and N50s ranging from 25 to 41 Kb, with an average of 1.6x genome coverage of ultra-long reads (>50 Kb) per sample (**Supplementary Table 3**). These longer-read N50s, paired with comparatively good quality scores, likely make these reads better suited for detecting large, complex structural variants compared to read N50s of 10 - 20 Kb, as they trade off read length for marginal gains in basecalling accuracy (*125*). The extended contiguity of our nanopore reads was particularly critical for spanning repetitive regions, such as transposable elements, which accounted for 54.5% of all SVs detected. A final caveat is that highly repetitive regions provide a challenge for accurate assembly and read mapping as we demonstrated with the MCH region in this study. While we validated our results by improving the assembly quality of this region, caution is warranted for SV calls in challenging-to-assemble regions of the genome, such as areas with dense tandem arrays, rapid evolution, and the presence of intergenic repeats in nearby regions (*126–128*). The variation graph confirmed 96.8% of our SV calls, demonstrating the robustness of our findings. However, a detailed physical map of the MHC and olfactory receptor loci would ultimately be required to definitively resolve their structures and confirm these enrichment patterns.

In this study, we provide a haplotype-resolved, near telomere-to-telomere genome assembly for *Boiga irregularis*. This assembly represents a critical resource for understanding the genomic basis of its remarkable invasion success in Guam. Our findings reveal that *B. irregularis* expanded dramatically on Guam despite extreme inbreeding, which is likely associated with a severe founder effect. Despite the bottleneck experienced by *B. irregularis*, we demonstrate that genetic diversity in the form of thousands of genomic structural variants was either maintained or re-established in chromosomal regions that harbor immune, stress response, and olfaction genes. These results highlight how using a more locus-specific approach that includes assessing multiple variant types can be implemented when measuring genetic diversity within a population. Accurately evaluating genetic diversity and adaptive potential in conservation genomics, where an overreliance on genome-wide summary measures of SNV-focused genetic diversity might lead to misdiagnosing a population’s or species’ viability, and concomitantly, a need for genetic rescue. The degree of inbreeding in *B. irregularis*, coupled with its putative adaptability and invasion success, provides a clear example of how bottlenecked species can escape the extinction vortex. We also discovered a substantial degree of sex chromosome heteromorphism in this species, raising questions about the evolutionary dynamics of sex chromosomes in snakes, particularly in the context of extreme inbreeding. The fully phased sex chromosomes of *B. irregularis* produced here may facilitate future studies of sex chromosome evolution in squamates. Finally, the availability of this high-quality assembly can aid future research exploring the genomic basis of invasion success in this species and represents a considerable advancement in genomic resources for developing management strategies for this highly detrimental invader and injurious species.

## METHODS

### Genome Sequencing, Assembly, and Phasing

A single female *Boiga irregularis* specimen was collected from Cocos Island, also known as Island Dånó, in 2021 (collection coordinates listed in **Supplementary Table 4**) to produce a genome assembly. The specimen was euthanized, and ∼100-300 μL blood was preserved in 1 mL DNA/RNA Shield (Zymo Research, Inc., Irvine, California, USA) and transferred from Guam on wet ice to the University at Buffalo, Buffalo, New York, USA. High-molecular-weight (HMW) DNA was extracted using Qiagen Genomic-tip (Qiagen, Inc., Hilden, Germany) kits following the manufacturer’s protocol. DNA was quantified on the Qubit 3.0 and checked for protein and RNA contamination using a NanoDrop 2000/2000c Spectrophotometer. A Short Read Eliminator Kit (Pacific Biosciences, Inc., Menlo Park, California, USA) was used to reduce the number of DNA fragments shorter than 25 Kb. Filtered HMW DNA was used in LSK114 ligation preparation and sequenced using a single R10.4.1 PromethION flow cell on the PromethION P24 instrument. This produced 74.5 Gb (∼42.6x coverage) of passed reads with a read-length N50 of 27 Kb. ONT long-reads were basecalled using the Super–accurate basecalling mode in Guppy v6.5.7 and filtered to retain only reads with basecalling accuracy of Q10 or higher and a minimum read length of 1 Kb. Approximately 46.0 Gb; ∼27.0x coverage Hi-C data were produced on the Illumina NovaSeq 6000 sequencing platform using DNA extracted from the blood of the same individual used for the genome assembly.

Oxford Nanopore Technologies (ONT) reads were corrected using the deep learning-based haplotype-aware read correction tool HERRO (*129*). Corrected long-reads and Hi-C data were then used as inputs in Hifiasm v0.19.6 (*67*) to produce a draft haplotype-resolved assembly. HERRO-corrected reads >30 Kb in length were trimmed or split using SeqKit2 v2.9.0 (*130*) to avoid known issues with contained read exclusion when using corrected ONT reads in Hifiasm (*131*). Hi-C reads were mapped to each contig-level assembly using bwa-mem2 v2.2 (*132*), then used to produce chromosome-level scaffolds using YaHS v1.2.2 (*133*). Finally, Juicebox (*134*) was used to visualize and manually correct misassemblies. The completeness of each assembled haplotype was assessed using BUSCO_v5 (*135*) performed online via gVolante2 (*136*) using the vertebrata_obd10 database.

### Telomere Identification

We used tidk v0.2.63 (*137*) to search for strict “AACCCT” tandem repeats (*137*) to characterize telomeric repeats in each assembled haplotype. A window size of 20 Kb and filtering for >21 consecutive tandem repeats were used to minimize noise from alternative repeat classes across the genome interfering with telomere classification.

### Genome Annotation

The Krabbenhoft Lab Annotation pipeline (https://github.com/KrabbenhoftLab/genome_annotation_pipeline) was used to produce gene annotations for each assembled haplotype. First, RepeatModeler v2.01 (*138*) was used along with the “Vertebrata” database from *Dfam_3* (*139*) to identify repetitive elements in each assembly. RepeatMasker v4.1.1 (*140*) was then used to identify and soft-mask repetitive elements using the custom repeat library generated for each haplotype. GeMoMa v1.71 (*141*) was then used to generate homology-based gene predictions for each repeat-masked haplotype using annotations from chromosome-level genome assemblies from sixteen snake species (**Supplementary Table 1**), which were downloaded from the Snake Multiomic Database (http://herpmuseum.cib.ac.cn/snake/), from ref. (*88*). Each assembly was used as a “reference” to predict gene models using the module GeMoMapipeline. Gene models generated from each reference species were aggregated and filtered to retain only those with the highest support values using the GeMoMa annotation finalizer using default parameters (*140*). For each haplotype, gene models were also generated using BRAKER3 v3.0.3 (*142*) with the same reference data used as input for GeMoMa predictions. Finally, EVidenceModeler v2.0.0 (*143*) was used to merge gene models produced from GeMoMa and BRAKER3 pipelines to produce a final genome annotations for each haplotype. Annotation completeness was assessed using BUSCO_v5 (*135*) online via gVolante2 (*136*) using the vertebrata_obd10 database.to produce gene annotations for each assembled haplotype. First, RepeatModeler v2.01 (*138*) was used along with the “Vertebrata” database from *Dfam_3* (*139*) to identify repetitive elements in each assembly. RepeatMasker v4.1.1 (*140*) was then used to identify and soft-mask repetitive elements using the custom repeat library generated for each haplotype. GeMoMa v1.71 (*141*) was then used to generate homology-based gene predictions for each repeat-masked haplotype using annotations from chromosome-level genome assemblies from sixteen snake species (**Supplementary Table 1**), from ref. (*88*). Each assembly was used as a “reference” to predict gene models using the module GeMoMapipeline. Gene models generated from each reference species were aggregated and filtered to retain only those with the highest support values using the GeMoMa annotation finalizer using default parameters (*140*). For each haplotype, gene models were also generated using BRAKER3 v3.0.3 (*142*) with the same reference data used as input for GeMoMa predictions. Finally, EVidenceModeler v2.0.0 (*143*) was used to merge gene models produced from GeMoMa and BRAKER3 pipelines to produce final genome annotations for each haplotype. Annotation completeness was assessed using BUSCO_v5 (*135*) online via gVolante2 (*136*) using the vertebrata_obd10 database.

### Long-Read Whole-Genome Resequencing

Eight additional *B. irregularis* samples were collected from Guam (collection coordinates listed in **Supplementary Table 4**) to produce whole-genome long-read resequencing data (also known as Population-Scale Long Read Sequencing, PLRS). The same methods for euthanasia, blood collection, and HMW DNA extraction used for the genome assembly sample were used for the resequencing samples. Long-read sequences were generated using the Oxford Nanopore Technologies (ONT) Native Barcode kit (SQK_NBD114.21) using four R10.4.1 flow cells on a PromethION P2 Solo instrument. This generated a total of 102.5 Gb of long-read data (range 9.0 – 19.8 Gb and 5.5 – 12.0x genome coverage per sample). The read N50s ranged from 26 to 41 Kb with an average of 33 Kb. Samples were collected under animal care/collection permits FORT IACUC 2021-01 and UOG2305.

### Analyses of Demographic History

We reconstructed the long-term demographic history of *B. irregularis* by first generating 18.5 Gb (∼11x coverage) of 150 bp paired-end Illumina short-read sequencing data on the NovaSeq 6000 platform. We then mapped these reads to a repeat-masked version of the hapZ assembly of *B. irregularis* using bwa-mem2 v2.2 (*132*), and detected SNVs using bcftools *mpileup* and *call* functions. SNV data were then used as input for the demographic history reconstruction tool Pairwise Sequentially Markovian Coalescence (PSMC v0.6.5-r67; (*69*). A nuclear mutation rate of 7.26e-09 mutations/site/generation, following (*76*) and a generation time of five years (per ref. (*75*) were used to scale the PSMC plots via the psmc_plot.pl utility.

To ensure that recent demographic fluctuations were not artifacts of the PSMC model, specifically the spurious population peaks identified by Hilgers et al. (*70*), we performed a series of sensitivity analyses on the atomic time interval parameters (-*p*). We first tested a model with an increased resolution of the initial time window (*-p 2+2+25*2+4+6*). This yielded negligible changes in *N_e_* trajectories compared to our primary model (*-p 4+25*2+4+6*). Following the recommendation to further decompose the first window for some problematic datasets, we also evaluated a highly fragmented model that split the initial time window into four (*-p 1+1+1+1+25*2+4+6*). This parameterization led to model instability, yielding biologically unrealistic *N_e_* estimates while failing to significantly alter the observed peak. Consequently, we retained the primary model for final inference, as the observed demographic trends remained robust across these diagnostic tests (**Supplementary Fig. 1**).

To assess statistical confidence and variance in the inferred demographic history, we performed a genomic bootstrapping with the PSMC. The initial .psmcfa file, representing the binned heterozygous sites across the genome, was partitioned into non-overlapping segments using the splitfa utility. We then generated 100 independent bootstrap replicates by randomly sampling these genomic segments with replacement (bootstrapping) using the -b flag. Each replicate was performed with the same parameterization as in the primary analysis. The resulting *N_e_*estimates from all 100 iterations were co-plotted with the primary genomic estimate to visualize the variance cloud and assess the robustness of the observed demographic fluctuations.

We assessed changes in the recent demographic history of *B. irregularis* by first calling SNVs using the ONT long-read resequencing data and using these data as input for the program GONE2 (*72*). First, ONT reads for each sample were mapped to the hapZ assembly with minimap2 (*74*) using the *–lr:hqae* flag, as is appropriate when mapping high-accuracy long-reads, and using the *–secondary=no* command to prevent secondary read-mappings. The BAM files from each sample were then used as input for bcftools *mpileup* using the following flags designed to optimize mpileup for ONT reads: *-q 10 -Q10 -C 50*. Next, SNVs were detected in each sample using bcftools *call* with the *-m, -v*, and *-f GQ,GP* options. Low-quality SNVs were removed using bcftools *filter -I QUAL>20* && *FORMAT/GQ>20* && *FORMAT/DP>=3*, ensuring that only SNVs with a quality score >20, a genotyping quality score >20, and only SNVs supported by at least three reads were retained. Only biallelic SNVs were retained using bcftools *view -v snps -m2 -M2*. Variant call format files for each sample were then merged into a single multi-sample vcf using bcftools *merge*. Finally, bcftools *view -i ‘MAF>0.05’ ‘F_MISSING>0.25’* was used to retain only SNVs with a minor allele frequency of <0.05, and allow for <25% missing data per locus (missing from 2 samples), as this approach leads to more loci retained and concomitantly higher accuracies when using low to moderate coverage data (*144*, *145*). Before running GONE2, excluded SNVs from the Z chromosome to avoid artifacts caused by sex chromosomes having different recombination and far higher linkage than autosomes (*72*). We also excluded data from seven chromosomes in the *B. irregularis* genome that were shorter than 20 Mb, as this is the software’s minimum length cutoff (*72*). We then ran GONE2 using the merged and filtered vcf file as input with default parameters except for *-g* 3, which was set to indicate that our data were low – moderate coverage, and *-b* 0.01, which was to account for the mean basecalling error rate of our ONT reads. To produce bootstrap replicates of these results, we reran GONE2 50 times with randomly selected subsamples of 500,000 SNVs using *-s 500,000* while the remaining parameters were identical to those used for the primary analysis. Effective population size trajectories from both PSMC and GONE were visualized using custom R scripts.

### Identifying Runs of Homozygosity

Runs of homozygosity (ROH) were identified in all samples using PLINK v1.9 (*146*). Before ROH detection, the unfiltered multi-sample vcf file described above (prior to MAF and missing data filtering) was converted to a PLINK binary format file using PLINK *--bfile --make-bed,* and --*geno 0.25* was added to remove markers with a call rate below 75% (equivalent <25% missing data). Following the recommendations of Meyermans et al. (*147*), no minor allele frequency filters were applied to the data, and the following parameters were applied to define an ROH segment in PLINK: *--homozyg --homozyg-window-snp 50 --homozyg-gap 500 --homozyg-density 60 --homozyg-snp 50 --homozyg-kb 50 --homozyg-het 1*. The raw ROH segment data and summary statistics were exported for further analysis using custom R scripts.

### Detection of Genomic Structural Variants

We used NextSV3 v3.2.0 (*148*) to detect genomic structural variants using the ONT long-read data from each sample. NextSV3 employs a reciprocal design that combines two read mappers and two SV callers to detect SVs for each sample. We ran the command *nextsv3.py -a minimap2+winnowmap* for each sample, so that minimap2 (*149*) and Winnowmap2 (*150*) were both used to map ONT reads to the hapZ assembly, producing two BAM files for each sample. Both BAM files were used as input for Sniffles2 v2.2 (*151*) and cuteSV v2.1.2 (*152*), resulting in four initial vcf files per sample: one for every combination of the two mappers and two SV callers. We then used SURVIVOR v1.0.7 (*153*) *merge* to merge the four vcf files and only retained an SV if it was detected in all four initial vcf files for each sample. To be merged, we required that both the start and end positions of SVs be within 1 Kb of each other. We then combined the final vcf files for all samples into one multi-sample vcf file and merged redundant SVs using the same parameters in SURVIVOR *merge*. Finally, we used SVJedi-Graph v1.2.1 (*82*) to convert the resulting multi-sample vcf file into a *B. irregularis* variation graph and tested the accuracy of SVs detected by NextSV3 by performing read-to-graph genotyping with ONT long reads for each sample, also using SVJedi-Graph. Only SVs assigned the same genotype by NextSV3 and SVJedi-Graph were retained and used in further analyses.

### Gene Ontology Enrichment Analysis

To identify Gene Ontology (GO) terms that were overrepresented in ROH and SV-affected genes, we used the Ensembl BioMart tool (https://www.ensembl.org/biomart) to collect data for the Indian cobra (*Naja naja*, Nana_v5), Mainland tiger snake (*Notechis scutatus*, TS10Xv2-PRI), Eastern brown snake (*Pseudonaja textilis*, EBS10Xv2-PRI), and Tropical clawed frog (*Xenopus tropicalis*, UCB_Xtro_10.0) genome assemblies. For each species, the corresponding dataset was selected in BioMart, and the following attributes were downloaded: Gene stable ID, GO term accession, and GO term name. Annotations were used to assign functional categories to all genes in the *B. irregularis* genome. The *find_enrichment.py* script from GOATOOLS (*154*) was used to detect significantly enriched GO terms for genes affected by SVs, as well as those contained within ROH deserts and ROH Islands, relative to a background set of all genes in the genome using Fisher’s exact tests. Benjamini-Hochberg procedure was used to adjust for multiple comparisons.

### Evaluation of Assembly Quality and Validation of Structural Variants in the MHC Locus

We evaluated the assembly quality and contiguity of the major histocompatibility complex (MHC) in the original *B. irregularis* assembly using comparative genomic mapping against the highly contiguous *B. cyanea* reference assembly (*86*), using D-GENIES (*155*). This whole-genome alignment revealed that the segment of chromosome 2 containing the *B. irregularis* MHC region was fragmented, containing 11 contig breakpoints across approximately 40 Mb. This fragmentation appears to have been an artifact of Hi-C scaffolding, which failed to correctly resolve the region’s dense repeat content. Our original, unscaffolded *B. irregularis* contigs mapped neatly to the corresponding *B. cyanea* MHC region, with the entire *B. irregularis* MHC region well-assembled and contained within just three large, highly contiguous Hifiasm contigs (**Supplementary Figure 3a-b**). We used RagTag (*156*) to perform a reference-guided re-scaffolding of these three contigs, using the *B. cyanea* assembly as the structural backbone for this localized region. This resulted in an 18.0% (48.6 Mb) increase in the total length of *B. irregularis* chromosome 2 while reducing the number of scaffolding gaps by one.

To assess whether the improved assembly of the MHC region impacted our finding that this region was rich in SVs, we reran the NextSV3 pipeline described above to detect SVs in the newly assembled chromosome. We also evaluated read coverage and mapping quality across the region in nonoverlapping 50 Kb windows using samtools 1.16.1 (*157*).

### Identification and Quantification of Structural Variant Runs of Homozygosity

To characterize structural variant runs of homozygosity (SV ROH) and compare them to traditional SNV-based ROH metrics, we defined an SV ROH as any genomic tract of 50 Kb or longer completely lacking heterozygous SVs, mirroring our methods for SNV ROH. We used *bcftools view -i ‘GT=“het”’* to filter the initial SV VCF file, isolating only heterozygous genotypes. This filtering step was executed independently for each sample to prevent removal of sites that were homozygous reference in one individual but variant in others. We employed bedtools v2.31.0 (*158*) *makewindows –w 50,000* to partition the reference genome into 50 Kb non-overlapping windows. We then used bedtools *intersect -c* to count the number of heterozygous SVs within each window for every individual sample. Windows containing zero heterozygous SVs were isolated and bedtools merge was used to combine adjacent heterozygous-free windows into continuous stretches, establishing the final SV ROH tracts for each sample. We calculated the cumulative length of SV ROH per chromosome for each sample, as well as the total number of base pairs structurally altered by heterozygous SVs. Statistical comparisons and data visualization were generated using custom R scripts.

### Sex Chromosome Identification and Size Validation

We used two approaches to identify the Z and W chromosomes in the *Boiga* assemblies. First, we used the synteny detection software GENESPACE v1.3.1 (*159*) to identify and visualize conserved synteny between *B. irregularis* haplotype assemblies and 16 other snake species with chromosome-level genome assemblies. All annotations were formatted and filtered to only the longest transcript using AGAT v0.9.2 (*160*); https://github.com/NBISweden/AGAT). For several of these species, the Z chromosome has previously been identified, including the common garter snake (*Thamnophis sirtalis* (*161*), which, like *Boiga,* is a member of the family Colubridae. Given that only one chromosome per assembly had conserved synteny with the Z chromosome in *P. guttatus,* we assumed that these chromosomes represent the Z chromosomes in the remaining assemblies, including *B. irregularis*. The W chromosome in *B. irregularis* was determined via retained synteny with the identified Z chromosomes, presumably representing pseudo-autosomal chromosome segments. Next, we mapped the ONT-reads the newly assembled *B. irregularis* assembly using minimap2 (*74*), which we determined to contain the Z chromosome, and used mosdepth v0.3.3 (*162*) to calculate and visualize regions of the genome with approximately one-half of the average read depth of the rest of the genome, following the approach used in previous work in snakes (*87*). This confirmed that the same chromosome with strongly conserved synteny with the Z chromosome in *P. guttatus* had approximately one-half the average genome-wide coverage in females, but not in males.

To validate the size difference between the putative Z and W chromosomes in *B. irregularis*, we mapped the putative W chromosome to the (larger) Z chromosome using minimap2 (*74*) and plotted the mapping locations and read-depth data from the long reads to check if regions of high read depth overlapped the regions of conserved synteny between the sex chromosomes. Indeed, regions of conserved synteny had read depths close to the genome-wide average while regions with approximately half the average read depth had no corresponding segment on the W chromosome. Thus, we conclude the low-coverage regions are those that have been lost via degeneration of the W chromosome in *B. irregularis*.

### Use of AI

Google Gemini 3 Flash model was used to produce human-readable annotations for the scripts that are being made publicly available with this manuscript. To accomplish this, we provided an unannotated version of each script to Gemini and used the following prompt: “Would you please provide some notes to this script that explain what each step of the pipeline is doing? Please don’t change or try to optimize the scripts otherwise. The goal is for the script’s purpose and function to be easily understood by someone unfamiliar with each method”. We then read and, when necessary, corrected the annotations and confirmed that none of the actual scripts were changed.

## Acknowledgements

We thank Marijoy Viernes, Christiana-Jo Quinata, Adam Perez, Jerilyn J. Calaor, Eben Paxton, and Corinna Pinzari for providing samples and/or assistance in processing and shipping tissues. Victor Albert, Charlotte Lindqvist, Omer Gokcumen, Dimitry Gorsky, Brendan Pinto, and members of the T. Krabbenhoft Lab provided helpful feedback on this research. Corinna Pinzari, Jonathan Richmond, and two anonymous reviewers provided constructive feedback that helped to improve this manuscript.

## Author Contributions

Conceptualization: CAO, LNG, TJK

Methodology: CAO, LNG, TJK

Software: CAO, SJF, SLC

Investigation: CAO, BMF, LNG

Formal Analysis: CAO, SJF, SLC

Resources: MGN, MRB, TJK

Data Curation: CAO, LNG, HMW

Writing – original draft: CAO, TJK

Writing – review & editing: CAO, SJF, HMW, SLC, MGN, MRB, LNG, TJK

Visualization: CAO

Supervision: TJK

Project Administration: TJK Funding Acquisition: TJK

## Data Availability

The *Boiga irregularis* genome assemblies have been deposited in NCBI with the hapZ and hapW haplotypes deposited under BioProject accessions PRJNA1332707 and PRJNA1333394, respectively. All raw sequencing data, including Oxford Nanopore long reads, Illumina short reads, and Hi-C data, are available in the NCBI Sequence Read Archive (SRA) under BioProject PRJNA1332707. The variation graph and gene model files are available at Dryad (https://doi.org/10.5061/dryad.g1jwstr6m). All scripts used for data processing and analysis are publicly available at https://github.com/KrabbenhoftLab/Boiga_irregularis_genome. All data are in the main text or the supplementary materials. The AI tool Google Gemini 2.5 Flash was used to annotate scripts for this project, and all code was reviewed, tested, and validated by the authors to ensure correctness and reproducibility.

## Ethics Statement

The collection and transfer of biological material was conducted under a Special Permit for Scientific Collection/Research License Nos. SC-21-004 and SCR-25-012 issued to Licensee Melia Nafus and Adam A. Perez, respectively. The exportation of samples from Guam was authorized under Certificate of Origin Permit No. CO-25-024. Any use of trade, firm, or product names is for descriptive purposes only and does not imply endorsement by the U.S. Government

